# A developmental stage- and Kidins220/ARMS-dependent switch in astrocyte responsiveness to brain-derived neurotrophic factor

**DOI:** 10.1101/2021.02.10.430582

**Authors:** Fanny Jaudon, Martina Albini, Stefano Ferroni, Fabio Benfenati, Fabrizia Cesca

## Abstract

Astroglial cells are key to maintain nervous system homeostasis, as they are able to perceive a wide variety of extracellular signals and to transduce them into responses that may be protective or disruptive toward neighboring neurons through the activation of distinct signaling pathways. Neurotrophins are a family of growth factors known for their pleiotropic effects on neuronal survival, maturation and plasticity. In this work, we investigated: (i) the signaling competence of embryonic and postnatal primary cortical astrocytes exposed to brain-derived neurotrophic factor (BDNF); and (ii) the role of the scaffold protein Kinase D interacting substrate/ankyrin repeat-rich membrane spanning (Kidins220/ARMS), a transmembrane protein that mediates neurotrophin signaling in neurons, in the astrocyte response to BDNF. We found a shift from a kinase-based response in embryonic cells to a predominantly [Ca^2+^]_i_-based response in postnatal cultures associated with the decreased expression of the full-length BDNF receptor TrkB. We also found that Kidins220/ARMS contributes to the BDNF-activated intracellular signaling in astrocytes by potentiating both kinase and [Ca^2+^]_i_ pathways. Finally, Kidins220/ARMS contributes to astrocytes’ homeostatic function by controlling the expression of the inwardly rectifying potassium channel (Kir) 4.1. Overall, our data contribute to the understanding of the complex role played by astrocytes within the central nervous system and identify Kidins220/ARMS as a novel actor in the increasing number of pathologies characterized by astrocytes’ dysfunctions.

## INTRODUCTION

Astroglia, the most abundant cell population of the central nervous system (CNS), is taking the stage in the neuroscience field because of its multifaceted role in the modulation of neural physiology at increasing levels of complexity, from the single synapse to higher circuits *(1)*. Key to astrocyte performances is the capability to sense a wide range of extracellular signals and consequently activate intracellular signaling pathways, which can be kinase-based, as for the ubiquitous mitogen-activated protein kinase (MAPK) (SHP2/Ras/ERK), signal transducer and activator of transcription (STAT) 1/3 and nuclear factor κB (NFκB) cascades *(2)*, or initiated via second messengers that trigger the release of Ca^2+^ ions from the endoplasmic reticulum *(3)*. Intracellular Ca^2+^ transients ([Ca^2+^]_i_) underlie the vast majority of astrocytes’ responses by mediating the release of a plethora of soluble factors globally referred to as gliotransmitters, which include neurotransmitters, trophic factors, hormones, cytokines and several other bio-active molecules. Gliotransmitters exert both autocrine and paracrine functions, thus controlling and coordinating the activity of astroglia itself and of neighboring neurons *(4)*.

While the mechanisms underlying astrocyte uptake/release of glutamate, ATP and other gliotransmitters, as well as the ability to respond to neurotransmitters such as glutamate, are well described and agreed upon by the scientific community, to what extent astrocytes produce, sense and respond to neurotrophins, such as nerve-growth factor (NGF) and brain-derived neurotrophic factor (BDNF), is still object of debate. Neurotrophins are mostly studied for their pleiotropic effects on neurons, as they control virtually every aspect of neuronal physiology, including maturation, axonal and dendritic differentiation, synaptogenesis and various forms of synaptic plasticity *(5)*. Besides their well-known effects on neurons, however, neurotrophins also modulate important aspects of astrocyte physiology *(6)*. Within the CNS, the most abundant neurotrophin is BDNF, which signals through its tropomyosin-related kinase B full-length (TrkB) and truncated (TrkB-T1) receptors *(5)*. Astrocytes predominantly express the truncated form of the BDNF receptor, which triggers release of Ca^2+^ from intracellular stores *(7)* and activates various kinase pathways *(8, 9)* independently of the full-length protein. The BDNF/TkB-T1 system modulates several astrocyte properties, such as astrocyte morphological maturation and their capability of sensing neurotransmission by controlling the membrane targeting of glycine and GABA transporters *(9–12)*. An aberrant activation of this pathway has been described in the experimental autoimmune encephalomyelitis rodent model for multiple sclerosis, where overexpressed TrkB-T1 induces neuronal death by promoting the release of nitric oxide *(13)*. Moreover, astrocytic TrkB-T1 contributes to neuropathic pain and neurological dysfunctions in rodent models of spinal cord injury and of amyotrophic lateral sclerosis *(14, 15)*. TrkB-T1 is arguably the predominant receptor isoform expressed in astrocytes, however some reports indicated that cultured astrocytes can also express full-length TrkB *(16, 17)*. Interestingly, some evidence indicates that TrkB expression by astrocytes occurs upon CNS injury or in the presence of chronic and inflammatory diseases, such as multiple sclerosis *(18–20)*. Importantly, the competence of astrocytes to perceive and respond to BDNF seems to be different in various brain areas *(21)*, thus adding a further layer of complexity to the system. Besides sensing extracellular BDNF through specific receptors, astrocytes also uptake and recycle proBDNF and BDNF *in vitro* and *in vivo (22, 23)*. An increasing body of evidence suggests that cultured astrocytes produce and secrete BDNF under basal conditions *(24, 25)*, and that BDNF expression is increased by specific stimuli *(26–30)*.

Although the contribution of astrocytes to neuronal function is becoming clear, the cellular and molecular mechanisms underlying the regulation of astrocyte-neuron communication mediated by neurotrophins remains incompletely understood. Kinase D interacting substrate/ankyrin repeat-rich membrane spanning (Kidins220/ARMS, henceforth referred to as Kidins220) is one of the several proteins involved in the neurotrophin pathways. Kidins220 is a four-pass transmembrane protein enriched in the nervous system, where it is expressed by both neurons and astrocytes and is involved in various aspects of neuronal differentiation and plasticity *(31, 32)*. Recently, we have shown that Kidins220 participates to the molecular events inducing spontaneous and evoked [Ca^2+^]_i_ transients in primary astrocytes, controlling the vulnerability of astrocytes to genotoxic stress and the maturation of co-cultured neurons *(33)*.

In this work, we have investigated the responsivity of astrocytes to BDNF and the role played by Kidins220 in astrocytic BDNF signaling. We first compared the signaling competence of embryonic and postnatal primary cortical astrocytes exposed to BDNF and found a developmental shift from a kinase-based response in embryonic cells to a predominantly [Ca^2+^]_i_-based response in postnatal cultures, which was paralleled by the reduced expression of full-length TrkB. Second, we addressed the role of Kidins220 in BDNF-induced signaling in astrocytes, and showed that Kidins220 significantly contributes to both kinase and [Ca^2+^]_i_ pathways. The data contribute to the understanding of the complex response of astrocytes to neurotrophins, describing a developmental stage- and Kidins220-dependent switch in astrocyte responsiveness to BDNF.

## RESULTS

### BDNF induces the activation of intracellular signaling pathways mediated by full-length TrkB and Kidins220 in embryonic astrocytes

To determine the role of Kidins220 in the modulation of BDNF signaling in astrocytes, the expression of TrkB was compared by western blot analysis in wild type and Kidins220^−/−^ primary astrocytes derived from E18.5 embryos at 15 days *in vitro* (DIV), a stage at which cultures have reached confluence. Kidins220^−/−^ astrocytes displayed a significant reduction of both full length and truncated forms of TrkB of around 40% and 50% respectively, compared to wild type cells, while the ratio between these two isoforms remained unchanged (**Fig. 1A**). We then compared the expression level of the main downstream effectors of the BDNF/TrkB pathway, such as mitogen-activated protein kinase (MAPK)1/2, phospholipase Cγ (PLCγ) and Akt. Whereas total protein levels of MAPK1/2 and Akt were unaffected, we found a significant reduction of PLCγ in Kidins220^−/−^ astrocytes (**Fig. 1B**). We then asked whether Kidins220 ablation in these cells affects the response to BDNF and TrkB-mediated signaling. Fifteen DIV astrocytes were treated with 50 ng/ml BDNF for 5 or 30 min and the phosphorylation levels of MAPK1/2, PLCγ and Akt were quantified by western blot with phospho-specific antibodies (**Fig. 1C**). While the basal phosphorylation levels of MAPK1/2 and Akt were unchanged in Kidins220^−/−^ astrocytes, we observed a reduction in their activation upon BDNF stimulation that reached statistical significance for MAPK1/2. We also detected a significant reduction of the basal phosphorylation level of PLCγ, while the percent increase in PLCγ phosphorylation induced by the BDNF was similar in wild type and Kidins220^−/−^ astrocytes. A similar trend was observed when lower concentrations of BDNF (1 ng/ml) were used; in this case, however, the fold change increase in MAPK phosphorylation was reduced compared to that observed with 50 ng/ml BDNF (**Suppl. Fig. 1**). Of note, we could not detect reliable phosphorylation of PLCγ under these experimental conditions (not shown). While astrocytes predominantly express the truncated form of TrkB, some reports indicate that primary astrocytes can also express full-length neurotrophin receptors under physiological conditions and/or upon specific stimuli *(17, 34, 35)*. Although, under our culturing conditions, primary glial cultures are predominantly represented by astrocytes with only a small percentage of microglial cells *(33)*, we wanted to confirm that the observed activation of signaling pathways was ascribable to astrocytes. Thus, we performed immunostaining experiments using anti-phospho TrkB (Tyr516, the phosphorylation site required for MAPK activation) and anti-GFAP antibodies. Indeed, stimulation with BDNF increased pTrkB fluorescence intensity in GFAP-positive cells, confirming that full length TrkB activation occurred in astrocytes (**Fig. 1D**). Altogether these results show that, similar to its function in neurons *(36, 37)*, Kidins220 is a crucial regulator of BDNF-dependent TrkB signaling also in astrocytes.

**Figure 1.**
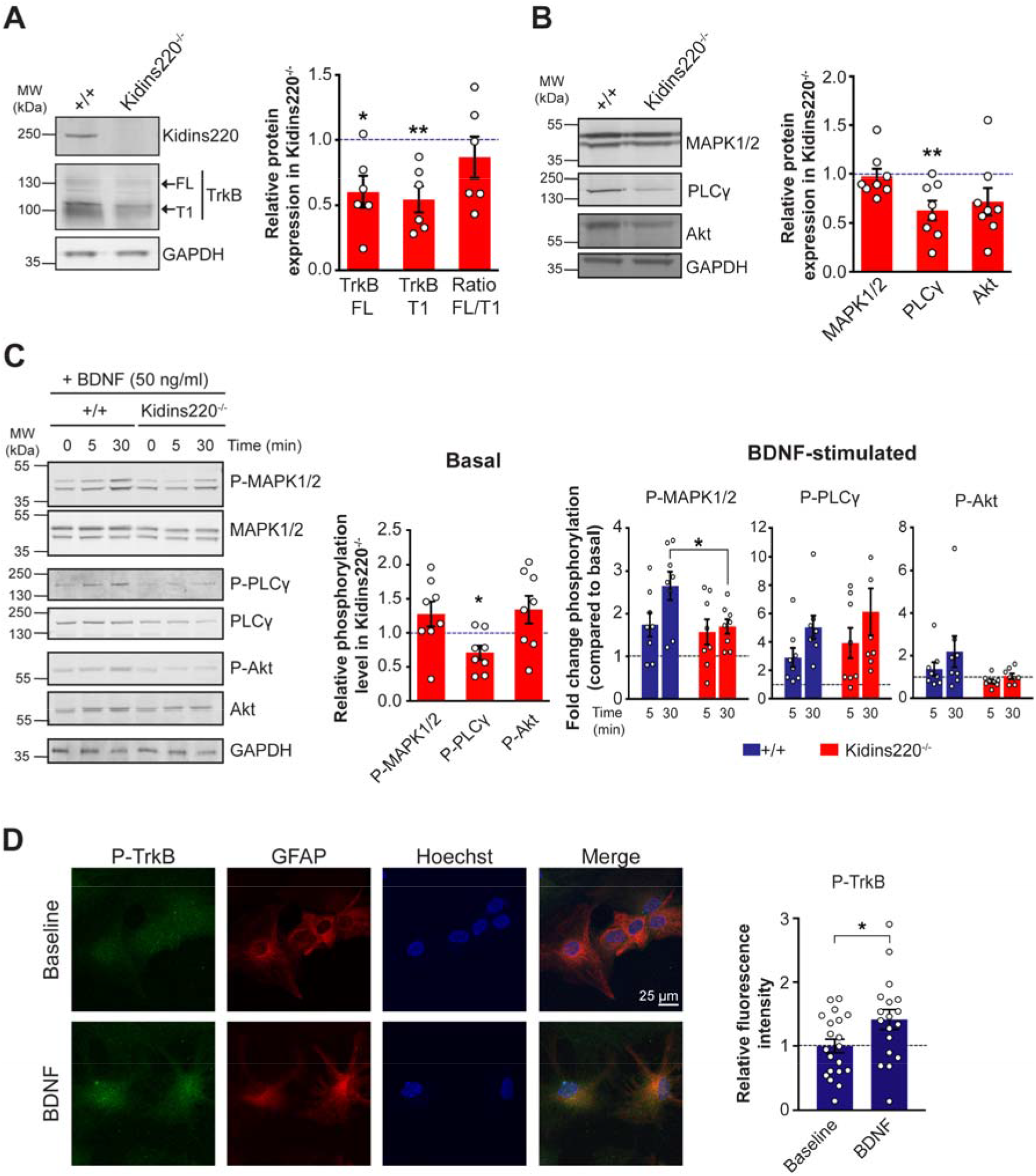
TrkB-dependent BDNF signaling is impaired in Kidins220^−/−^ embryonic astrocytes. (**A,B**) Protein extracts from wild type (+/+) and Kidins220^−/−^ embryonic astrocyte cultures at 15 DIV were analyzed by western blotting using anti-Kidins220 and anti-TrkB antibodies (**A**); anti-MAPK1/2, anti-PLCγ and anti-Akt antibodies (**B**). Representative immunoblots are shown on the left; quantification of immunoreactive bands is on the right. The intensity of bands from Kidins220^−/−^ samples was normalized to the corresponding bands from wild type samples within the same nitrocellulose membrane. *p<0.05, **p<0.01, one sample Student’s *t*-test, n=6 (A) and n=8 (B) wild type and Kidins220^−/−^ cultures. (**C**) Wild type and Kidins220^−/−^ astrocyte cultures were treated with 50 ng/ml BDNF for 5 and 30 min or left untreated (time 0). Lysates were analyzed for phosphorylated MAPK1/2 (Thr202/Tyr204), PLCγ (Tyr783) and Akt (Ser473). Membranes were subsequently stripped and re-probed for the total amount of the same protein. *Left:* Representative immunoblots. *Middle:* Basal phosphorylation levels of MAPK1/2, PLCγ and Akt in untreated wild type and Kidins220^−/−^ lysates. Data were analyzed as in (A,B). *Right:* Time dependence of MAPK1/2, PLCγ and Akt phosphorylation upon BDNF stimulation in wild type and Kidins220^−/−^ astrocytes. The graphs express the fold change activation of MAPK1/2, PLCγ and Akt compared to the untreated phosphorylation levels for each genotype, set to 1 (dashed line in all graphs). The fold change activation was calculated first by making the ratio of the intensity of phosphorylated bands to the total amount of protein for each sample then by dividing the phospho:total ratio of the treated samples by the phospho:total ratio of untreated control samples. For MAPK, we report the sum of MAPK1 and MAPK2 immunoreactivity. *p<0.05, unpaired Student’s *t*-test, n=8 for both wild type and Kidins220^−/−^ cultures. In all experiments, GAPDH was used as a loading control. (**D**) *Left:* Representative confocal images of untreated and BDNF-treated (1 ng/ml for 5 min) wild type astrocytes stained with anti-pTrkB (Tyr516, green) and anti-GFAP (red) antibodies, and with Hoechst to visualize nuclei. Scale bars, 25 μm. *Right:* quantification of pTrkB fluorescence intensity in GFAP positive cells. *p<0.05, unpaired Student’s *t*-test, n=20 and 18 fields for untreated and BDNF-treated cells, respectively, from three independent preparations. Data are reported as the average pTrkB fluorescence intensity of all GFAP+ cells in each field. Values are expressed as mean ± S.E.M. in all panels.

### The expression of full-length TrkB and the activation of BDNF-dependent kinase pathways are reduced in wild-type postnatal astrocytes

Astrocytes undergo important physiological changes from the embryonic development to the postnatal period, accompanied by variations in the expression pattern of several signaling proteins *(38)*. In order to study the developmental changes associated to the BDNF-TrkB system, we prepared primary astrocyte cultures from P0-P1 pups from Kidins220^+/lox^ X Kidins220^+/lox^ breeding couples, which allowed us to have both Kidins220^+/+^ and Kidins220^lox/lox^ animals in the same litter. We chose this strategy since the full knockout of Kidins220 causes embryonic lethality, thus precluding the possibility of performing postnatal dissection *(37, 39)*. We first compared the expression levels of full-length and truncated TrkB receptors in wild type embryonic and postnatal preparations and found a specific reduction of the full-length isoform in postnatal cultures, leading to a decreased full-length/truncated ratio compared to embryonic cells (**Fig. 2A**). Exposure of wild type postnatal cultures to 50 ng/ml BDNF led to an appreciable increase in Akt and MAPK1/2 phosphorylation, although the fold-increase in MAPK1/2 phosphorylation was lower than that observed in embryonic cultures (1.2-fold versus 2.5-fold at 30 min, compare **Fig. 2B** and 1C), a likely consequence of the lower expression of the full-length TrkB receptor. Similar results were observed when a lower concentration of BDNF (1 ng/ml) was used (**Suppl. Fig. 2**).

**Figure 2.**
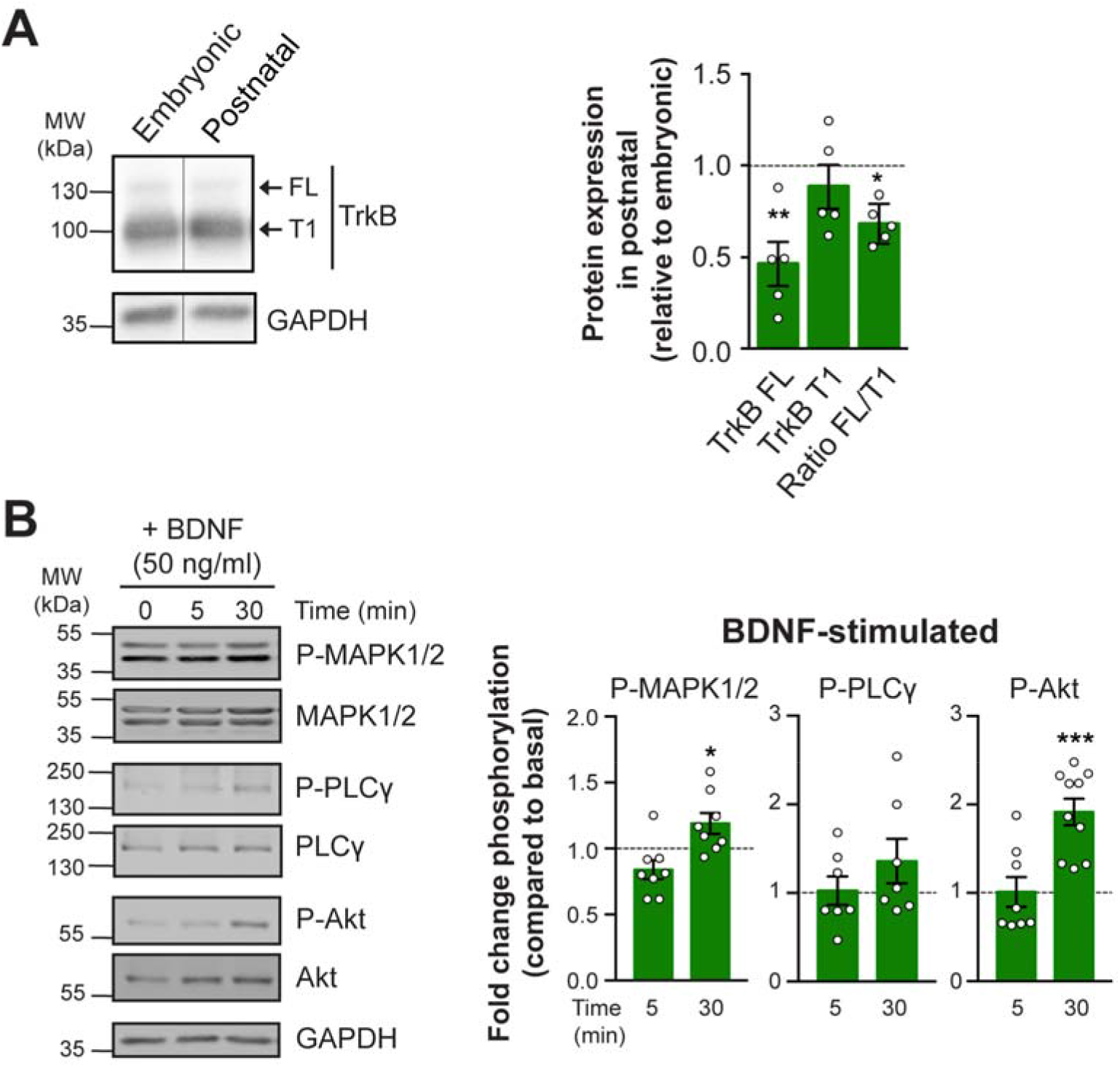
Expression of full-length TrkB is reduced in postnatal astrocytes. (**A**) Protein extracts from wild type embryonic and postnatal cultures were analyzed by western blotting with anti-TrkB antibodies. A representative immunoblot is shown in the left panel; quantification of immunoreactive bands is in the right panel. The intensity of bands from postnatal samples was normalized to the corresponding bands from wild type embryonic samples within the same nitrocellulose membrane. *p<0.05, **p<0.01, one sample Student’s *t*-test, n=5 wild type embryonic and postnatal cultures. (**B**) Postnatal wild type astrocyte cultures were treated with 50 ng/ml BDNF for 5 and 30 min or left untreated (time 0). Lysates were analyzed for phosphorylated MAPK1/2 (Thr202/Tyr204) and Akt (Ser473). Membranes were subsequently stripped and re-probed for the total amount of the same protein. *Left:* Representative immunoblots. *Right:* Time dependence of MAPK1/2 and Akt phosphorylation in response to BDNF in wild type and Kidins220^lox/lox^ astrocytes. The graphs express the fold change activation of MAPK1/2 and Akt compared to the untreated phosphorylation levels for each genotype, set to 1 (dashed line in all graphs). The fold change activation was calculated as described in Fig. 1. For MAPK, we report the sum of MAPK1 and MAPK2 immunoreactivity. *p<0.05, ***p<0.001, one sample Student’s *t*-test compared to baseline, n=7-8 independent cultures. Values are expressed as mean ± S.E.M. in all panels.

### Kidins220 controls the expression of both full-length and truncated TrkB receptors, but is dispensable for the activation BDNF-dependent kinase pathways in postnatal astrocytes

Primary Kidins220^lox/lox^ cultures were subsequently infected with lentiviruses expressing either the active form of the Cre recombinase to induce Kidins220 depletion, or a functionally inactive Cre (ΔCre) as control (*(40)*; see Materials and Methods for details). We refer to these cultures as ‘Kidins220-depleted’ (Kidins220^lox/lox-Cre^) and ‘control’ (Kidins220^lox/lox-ΔCre^). Cre-induced Kidins220 depletion in postnatal preparations led to a reduction of both full-length and truncated TrkB expression, with the full-length/truncated ratio remaining unaltered, a result in full agreement with that obtained on embryonic cultures (**Fig. 3A**). We then analyzed the signaling capability of postnatal cultures in the presence or absence of Kidins220. The basal expression levels of MAPK1/2, PLCγ and Akt were not affected by Kidins220 depletion (**Fig. 3B**). In contrast to the data obtained in embryonic cultures, removal of Kidins220 in postnatal cultures did not affect the activation of BDNF-induced MAPK1/2, PLCγ and Akt phosphorylation (**Fig. 3C** and **Suppl. Fig. 3**).

**Figure 3.**
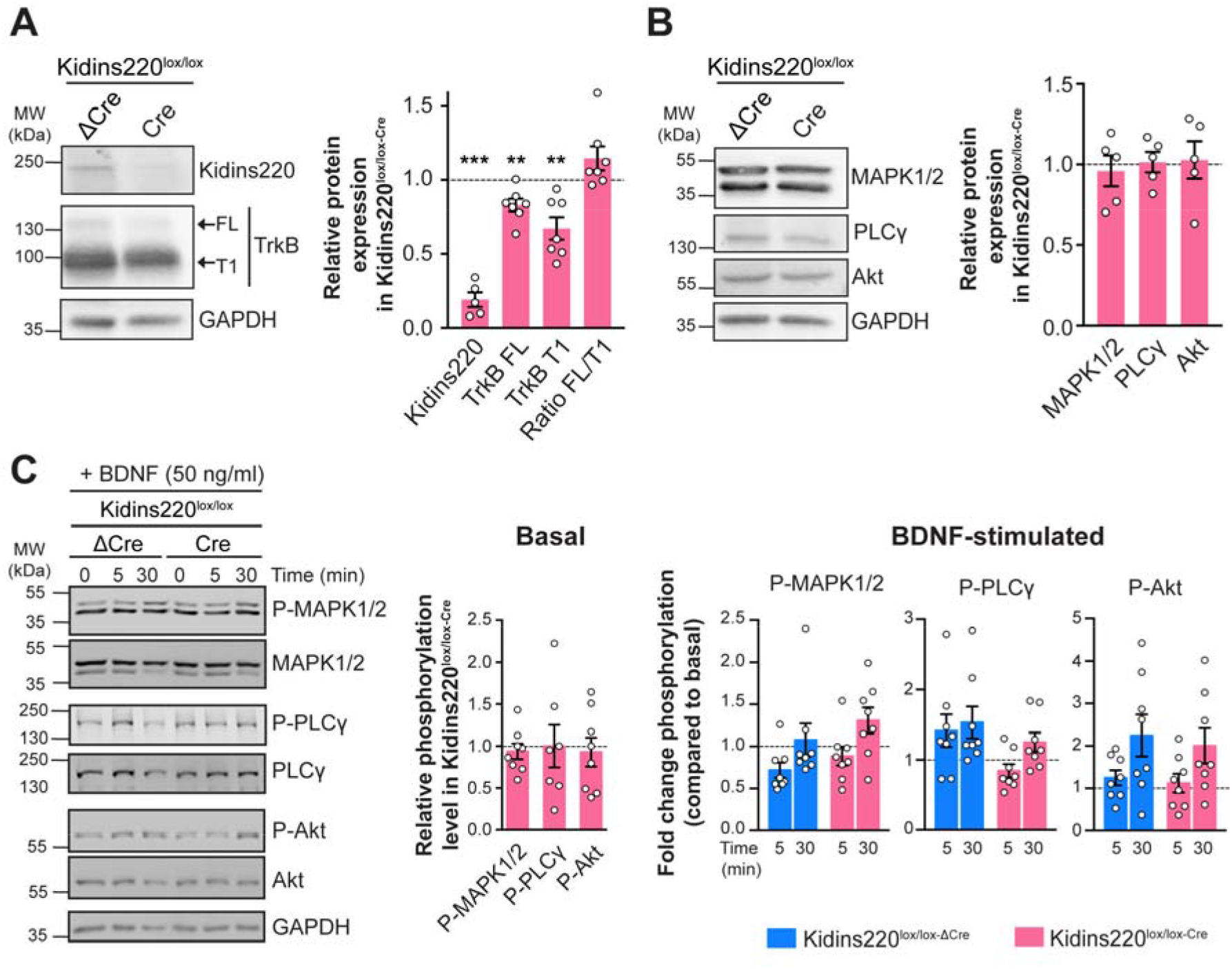
Removal of Kidins220 in postnatal astrocyte cultures does not alter TrkB-dependent BDNF signaling. (**A,B**) Protein extracts from Kidins220^lox/lox^ P0-P1 astrocyte cultures infected with lentiviruses encoding catalytically dead (ΔCre) or active Cre recombinase were analyzed by western blotting using anti-Kidins220 and anti-TrkB antibodies (**A**); anti-MAPK1/2, anti-PLCγ and anti-Akt antibodies (**B**). Representative immunoblots are shown on the left; quantification of immunoreactive bands is on the right. The intensity of bands from Kidins220^lox/lox-Cre^ samples was normalized to the corresponding bands from Kidins220^lox/lox-ΔCre^ samples within the same nitrocellulose membrane. **p<0.01, ***p<0.001, one sample Student’s *t*-test, n=5-7 Kidins220^lox/lox^ cultures. (**C**) Kidins220^lox/lox-Cre^ and Kidins220^lox/lox-ΔCre^ astrocytes were treated with 50 ng/ml BDNF for 5 and 30 min or left untreated (time 0). Lysates were analyzed for phosphorylated MAPK1/2 (Thr202/Tyr204) and Akt (Ser473). Membranes were subsequently stripped and re-probed for the total amount of the same protein. *Left:* Representative immunoblots. *Middle:* Basal phosphorylation levels of MAPK1/2 and Akt in untreated lysates. Data were analyzed as in (A,B). *Right:* Time dependence of MAPK1/2 and Akt phosphorylation upon BDNF stimulation in Kidins220^lox/lox-Cre^ and Kidins220^lox/lox-ΔCre^ astrocytes. The graphs express the fold change activation of MAPK1/2 and Akt compared to the untreated phosphorylation levels for each genotype, set to 1 (dashed line in all graphs). The fold change activation was calculated as described in Fig. 1. For MAPK, we report the sum of MAPK1 and MAPK2 immunoreactivity. p>0.05, unpaired Student’s *t*-test, n=8 independent Kidins220^lox/lox^ cultures. Values are expressed as mean ± S.E.M. in all panels.

Altogether, the data presented in Figure 2 and 3 show that (i) the expression of full-length TrkB and the activation of kinase pathways upon exposure to BDNF are reduced in wild type postnatal astrocytes compared to embryonic cells; and (ii) the expression of both full-length and truncated TrkB receptors is further reduced in the absence of Kidins220, however the limited activation of kinase pathways induced by BDNF is not impacted upon Kidins220 depletion.

### Depletion of Kidins220 impairs BDNF-induced [Ca^2+^]_i_ transients in postnatal astrocytes

We subsequently investigated the role played by Kidins220 in BDNF-induced [Ca^2+^]_i_ signaling by monitoring BDNF-induced [Ca^2+^]_i_ transients in astrocytes *(7)*. In embryonic cultures, very few cells responded to BDNF stimulation (9.8% for +/+ and 7.4% for Kidins220^−/−^, **Fig. 4A**) and responsive cells displayed modest [Ca^2+^]_i_ variations (**Fig. 4B-C**). In contrast, around 30-35% of cells in postnatal cultures responded to BDNF with sizable [Ca^2+^]_i_ peaks (**Fig. 4D-F**). Whereas the percentage of responding cells was not affected by Kidins220 depletion (**Fig. 4D**), the average amplitude of the BDNF-induced [Ca^2+^]_i_ transients was significantly reduced in absence of Kidins220 (**Fig. 4E** and **F**; # p < 0.05 genotype effect). To investigate the contribution of the different isoforms of TrkB in this process, we repeated the same experiments in the presence of either the tyrosine kinase blocker K-252a *(37)* to inhibit specifically the full-length TrkB isoform, or the TrkB antagonist ANA-12 *(41)* to block both full-length and truncated TrkB receptors. Pre-incubation of the cells with K-252a did not affect the percentage of BDNF-responding cells (**Fig. 4D**, shaded). However, the amplitude of the observed [Ca^2+^]_i_ influx was reduced independently of the presence of Kidins220 (**Fig. 4F** and **G**; genotype effect: p=0.0432 (#); K-252a treatment effect: p<0.0001 (***); treatment x genotype interaction: p=0.2197, see legend for more details), suggesting that the BDNF-induced [Ca^2+^]_i_ variations involved full-length TrkB receptors. BDNF-stimulation in presence of ANA-12 did not induce any response, confirming that the residual signal was attributable to TrkB-T1 activation (**Fig. 4H**). Furthermore, the observed response was completely dependent on PLCγ activity, as shown by the absence of BDNF-induced [Ca^2+^]_i_ transients in the presence of the specific PLCγ inhibitor U73122 (**Fig. 4I**).

**Figure 4.**
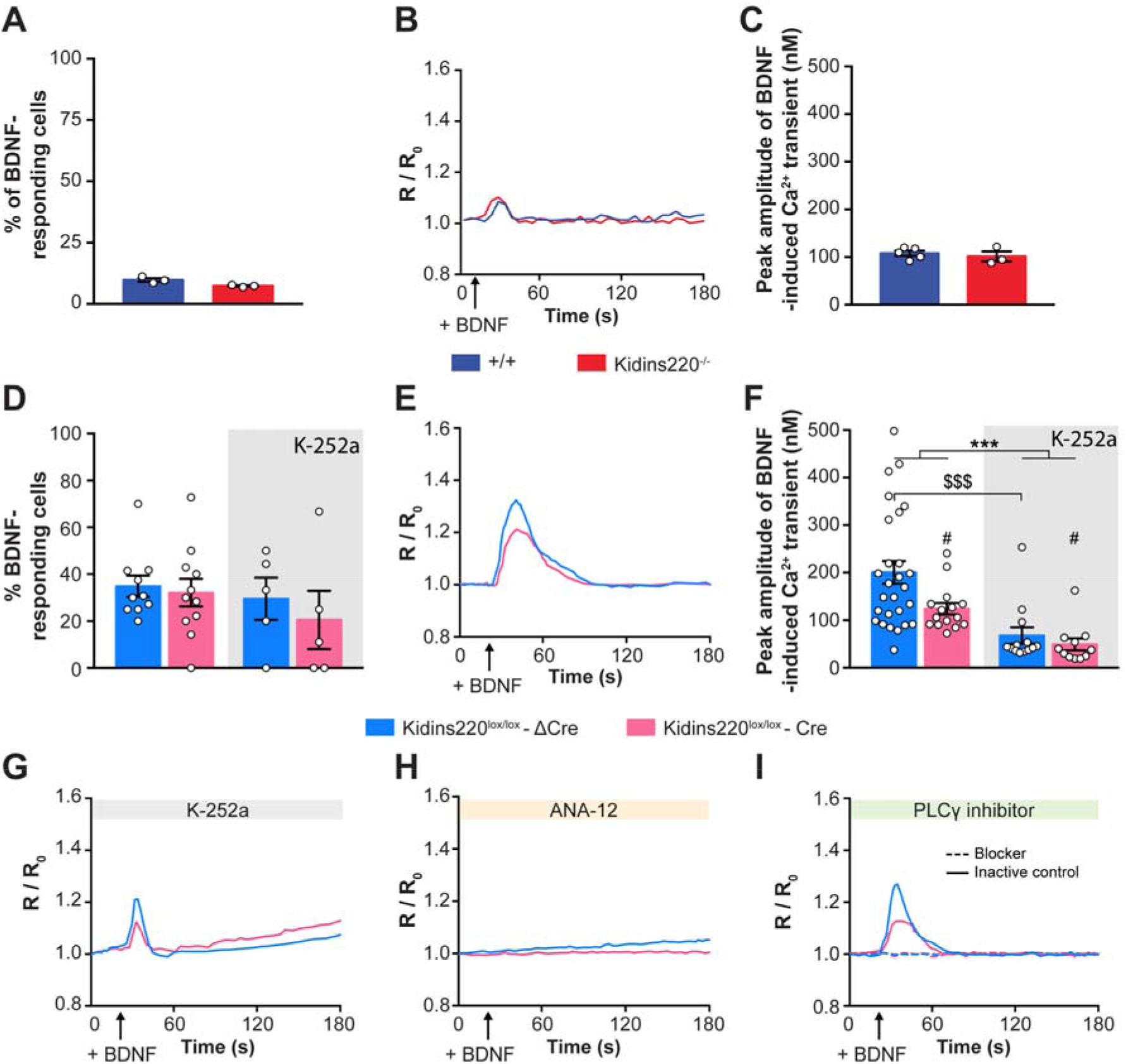
Depletion of Kidins220 in postnatal cultures impairs BDNF-induced calcium transients. (**A-C**) BDNF-induced [Ca^2+^]_i_ transients in embryonic wild type and Kidins220^−/−^ astrocytes. (**A**) Percentage of cells displaying Ca^2+^ transients in response to BDNF (20 ng/ml) stimulation. n=3 for both wild type and Kidins220^−/−^ cultures. [Ca^2+^]_i_ transients were defined when R/R_0_ ≥ 0.025. (**B,C**) Time-course of calcium transients (**B)** and peak amplitude (**C**) of BDNF-evoked Ca^2+^ levels in wild type and Kidins220^−/−^ astrocytes. (**D-I**) BDNF-induced [Ca^2+^]_i_ transients in postnatal Kidins220^lox/lox-Cre^ and Kidins220^lox/lox-ΔCre^ astrocytes. (**D**) Percentage of cells displaying Ca^2+^ transients in response to BDNF (20 ng/ml) stimulation in the presence or absence of the tyrosine kinase blocker K-252a. n=5 and 10 Kidins220^lox/lox^ cultures with and without K-252a, respectively. (**E,F**) Time-course of calcium transients (**E**) and peak amplitude (**F**) of BDNF-evoked Ca^2+^ influx in Kidins220^lox/lox-Cre^ and Kidins220^lox/lox-ΔCre^ astrocytes in the absence or presence of K-252a. Two-way ANOVA followed by Tukey’s multiple comparison test; ***p<0.001 versus untreated, #p<0.05 versus Kidins220^lox/lox-ΔCre^, $$$p<0.001 versus untreated Kidins220^lox/lox-ΔCre^ (genotype effect: F(1,62)=4.262, p=0.0432 (#); K-252a treatment effect: F(1, 62)=20.17, p<0.0001 (***); treatment x genotype interaction: F(1, 62)=1.538, p=0.2197), n=27 and 15 cells from 6 independent cultures for Kidins220^lox/lox-ΔCre^ and Kidins220^lox/lox-Cre^ cells in absence of K-252a, and n=13 and 11 from 6 independent cultures for Kidins220^lox/lox-ΔCre^ and Kidins220^lox/lox-Cre^ cells in presence of K-252a. (**G-I**) Time-course of calcium transients evoked by BDNF in Kidins220^lox/lox-ΔCre^ and Kidins220^lox/lox-Cre^ astrocytes in the presence of the TrkB receptor kinase inhibitor K-252a (**G**), the pan-TrkB antagonist ANA-12 (**H**), the active PLCγ inhibitor U73122 (dashed lines) or its inactive analogue U73343 (solid lines) (**I**). Values are expressed as mean ± S.E.M. in all panels.

Altogether, these data show that BDNF induce modest [Ca^2+^]_i_ events in embryonic astrocytes, while appreciable [Ca^2+^]_i_ responses are elicited in postnatal cells. Postnatal [Ca^2+^]_i_ transients are fully dependent on TrkB receptors, as shown by their complete inhibition in the presence of ANA-12. Interestingly, part of this response is mediated by the full-length receptor, as shown by the partial reduction of amplitude in the presence of K-252a. Moreover, our results show that Kidins220 plays an active role in this process, as in its absence the amplitude of [Ca^2+^]_i_ events is reduced.

### BDNF stimuli promote gene transcription in postnatal astrocyte cells

To shed light on the functional effects of BDNF signaling in astroglia, we analyzed the transcriptional profile of cultures treated with 1 ng/ml BDNF for three days. We focused on four genes whose expression is crucial for astrocytes physiology: the glutamate transporter 1 (GLT-1), the inwardly rectifying K^+^ channel 4.1 (Kir4.1), aquaporin 4 (Aqp4) and connexin 43 (Cnx43) (**Fig. 5**). While the expression of GLT-1, Kir4.1 or Cnx43 was not sensitive to BDNF, at least within the time frame of our experiment, Aqp4 transcription was selectively reduced upon chronic BDNF stimulation (* p<0.05, treatment effect). While Kidins220 deficiency did not alter the response to BDNF, it selectively and markedly reduced the expression of Kir4.1 independently of BDNF (## p<0.01, genotype effect).

**Figure 5.**
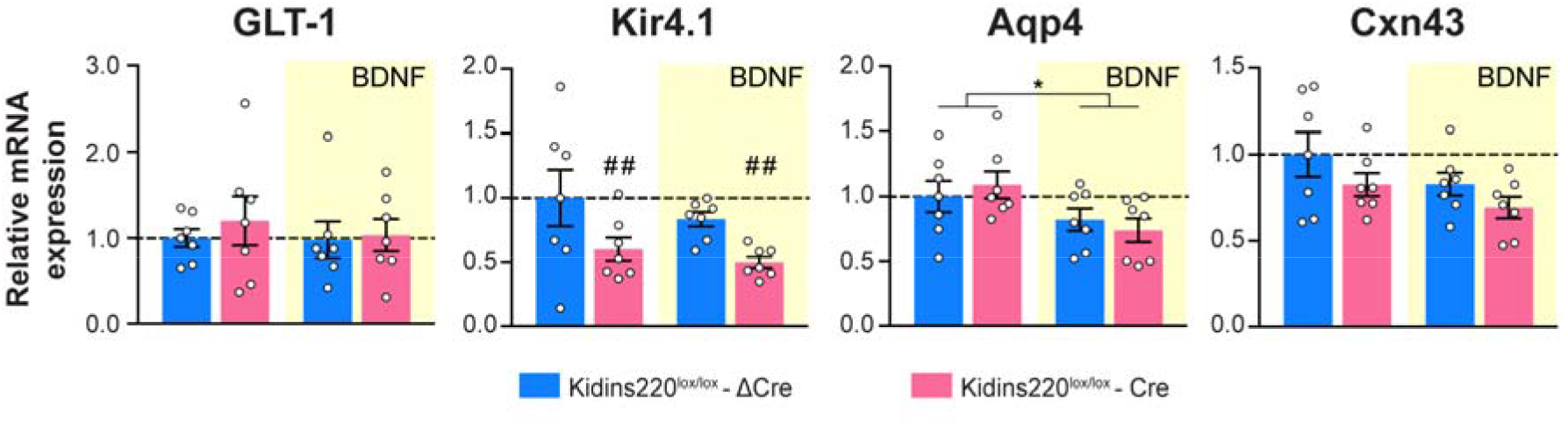
Transcription of Aqp4 and Kir4.1 in postnatal astrocytes are reduced by chronic BDNF and Kidins220 deficiency, respectively. mRNA expression profile of glutamate transporter 1 (GLT-1), inwardly rectifying potassium channel 4.1 (Kir4.1), aquaporin 4 (Aqp4) and connexin 43 (Cnx43) in postnatal Kidins220^lox/lox-ΔCre^ and Kidins220^lox/lox-Cre^ astrocytes with or without chronic BDNF treatment (1 ng/ml for 72 h, see Materials and Methods). mRNA expression in the various samples was normalized to the values of untreated Kidins220^lox/lox-ΔCre^ samples within the same RT-qPCR plate. Two-way ANOVA followed by Tukey’s multiple comparison test. *p<0.05 *versus* untreated, ##p<0.01 *versus* Kidins220^lox/lox-ΔCre^. **GLT-1:** Genotype effect: F(1, 24)=0.3813, p=0.5427; BDNF treatment effect: F(1, 24)=0.2042, p=0.6554; treatment x genotype interaction: F(1, 24)=0. 1262, p=0.7255. **Kir4.1:** Genotype effect: F(1, 24)=8.817, p=0.0067 (##); BDNF treatment effect: F(_1_,_24_)=1.179, p=0.2883; treatment x genotype interaction: F(1,24)=0.06038, p=0.8080. **Aqp4:** Genotype effect: F(1, 24)=0.002003, p=0.9647; BDNF treatment effect: F(1, 24)=6.817, p=0.0153 (*); treatment x genotype interaction: F(1, 24)=0.7202, p=0.4045. **Cnx43:** Genotype effect: F(1, 24)=3.302, p=0.0817; BDNF treatment effect: F(1, 24)=3.183, p=0.0870; treatment x genotype interaction: F(1, 24)=0.04637, p=0.8313. n=7 independent Kidins220^lox/lox^ cultures. Values are expressed as mean ± S.E.M. in all panels.

These data suggest that: (i) chronic exposure to BDNF alters the expression of specific astrocyte genes, and (ii) Kidins220 plays a role in the maintenance of astrocyte homeostasis and potassium buffering function by modulating the expression of Kir4.1.

## DISCUSSION

The main findings of this work can be summarized as follows: (i) primary cultured astrocytes express predominantly the TrkB-T1 isoform, however they do express a small amount of full-length TrkB receptor that is signaling competent, as shown by the BDNF-induced activation of the MAPK pathway, and whose expression declines in postnatal cells; (ii) astrocyte cultures show a stage-dependent shift in their response to BDNF, whereby embryonic cells mostly respond by activating intracellular phosphorylation pathways (MAPK1/2, PCLγ and Akt) and are not competent to induce sizable BDNF-dependent [Ca^2+^]_i_ transients, while postnatal cells respond to BDNF stimuli inducing robust variations of [Ca^2+^]_i_ and limited activation of intracellular signaling cascades; (iii) the scaffold protein Kidins220 participates to TrkB signaling both in embryonic and postnatal astrocytes and modulates the expression of fundamental astrocyte genes such as the inward rectifier potassium channel Kir 4.1. A schematic representation of our results is shown in **Fig. 6**.

**Figure 6.**
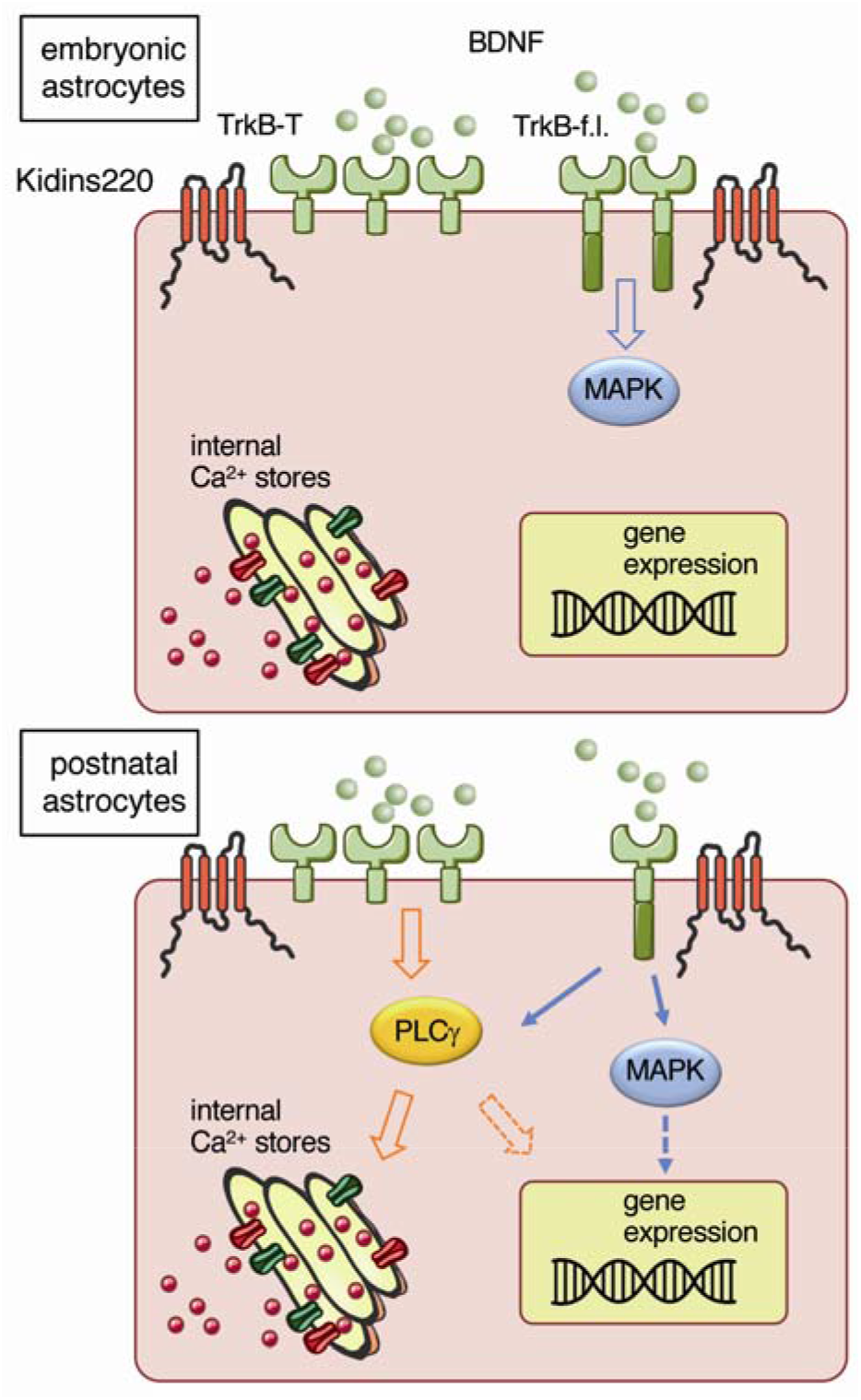
Kidins220 differentially contributes to BDNF signaling in astrocytes. We propose a model whereby embryonic astrocytes (top) mostly respond to BDNF stimuli through full-length TrkB-dependent activation of intracellular signaling cascades, while in postnatal cells (bottom) BDNF mainly activates intracellular [Ca^2+^]_i_ transients via PLCγ. Kidins220 contributes to the sustained activation of the MAPK pathway in embryonic cells, whereas it modulates Kidins220 BDNF-induced [Ca^2+^]_i_ transients in postnatal cells. Moreover, chronic BDNF treatment promotes the transcription of selected genes, such as Aqp4, in postnatal cells.

Extensive literature supports the notion that TrkB-T1 is the main isoform expressed in astroglia *(12, 42)*; however experimental evidence also points at a limited expression of the full-length receptor, at least *in vitro (16, 17)*. Moreover, reactive astrocytes express the full-length TrkB receptor upon CNS injury *(18)* and chronic diseases *(19, 20)*, raising the question of whether the astrocyte expression of full-length TrkB under pathological conditions recapitulates intracellular pathways that are normally active during development. Re-activation of developmental programs is indeed a known feature of reactive astrocytes *(43)* and of astrocyte-derived tumors such as gliomas *(44–46)*, and the BDNF/TrkB pathway is overexpressed in several brain tumors *(47)*. Consistent with previous observations *(17)*, our data show that the expression of full-length TrkB decreases from embryonic to postnatal astrocytes, supporting the idea that the intracellular pathways activated by this receptor could be active during embryogenesis and re-activated in the presence of neural pathologies. This finding is potentially relevant in that it shows that cultured astrocytes retain - to some extent - the identity of the tissue from which they were initially isolated *(48, 49)*.

For what concerns the signaling pathways activated by BDNF in astrocytes, our data indicate that activation of the canonical (MAPK1/2, PCLγ and Akt) phosphorylation pathways is predominant in embryonic cultures, while the same cascades are activated to a reduced extent in postnatal cells, in agreement with the reduced expression of full-length TrkB. Our embryonic and postnatal cultures are predominantly composed of astrocytes, with only a small (<10%) contamination of other glial cells, mostly microglia *(33)*. Although we cannot exclude that a small component of the activated pathways observed in western blot experiments is coming from cells other than astrocytes, the strong induction of protein phosphorylation observed upon BDNF treatment by western blotting, together with the increased phospho-TrkB staining in GFAP+ cells revealed by fluorescence microscopy analysis indicate that the observed changes in intracellular signaling pathways are predominantly attributable to astrocytes. The mechanisms underlying BDNF-induced [Ca^2+^]_i_ transients become fully functional in postnatal astrocytes; indeed, only modest BDNF-induced [Ca^2+^]_i_ elevations are observed in embryonic cultures. We hypothesize that such limited response could be ascribable to a still immature coupling between the Ca^2+^ receptors and channels on endoplasmic reticulum and on the plasma membrane, which together coordinate the store-operated Ca^2+^ entry (SOCE) mechanism in mature astrocytes *(3)*. The [Ca^2+^]_i_ transients observed in postnatal cells upon BDNF administration comprise two components. The first, which accounts for about 40% of the observed signal, is mediated by fulllength TrkB and is sensitive to pre-incubation with K-252a. The complete ablation of [Ca^2+^]_i_ transients observed in the presence of the pan-TrkB inhibitor ANA-12 reveals that the second and predominant component is dependent on TrkB-T1, as previously reported *(7)*. The complete dependence of [Ca^2+^]_i_ variations on PLCγ suggests that the pathways activated by both receptors converge on SOCE. While only 40% of postnatal astrocytes were responsive to BDNF in terms of [Ca^2+^]_i_ transients, the majority of GFAP+ cells were immunoreactive for full-length TrkB phosphorylation in embryonic cultures. Our calcium imaging data confirm that the capability of astrocytes to initiate BDNF-induced [Ca^2+^]_i_ transients varies depending on the degree of differentiation of the cells. Thus, we propose that the expression of full-length TrkB is a general feature of embryonic cells; whether BDNF-responsive astrocytes in postnatal cultures represent a specific subpopulation endowed with distinct physiological functions is a relevant issue that will be addressed in future investigations. Interestingly, chronic treatment with BDNF reduced Aqp4 expression, indicating that the neurotrophin is capable to induce long-term changes in astrocytes, potentially impacting on their physiology and on their capability to maintain nervous system homeostasis.

The scaffold protein Kidins220 is highly expressed in the nervous system during embryonic development and plays a fundamental role in modulating neuronal survival and maturation; indeed, its complete ablation leads to embryonic lethality with widespread apoptosis in the central and peripheral nervous system *(37, 39)*. Kidins220 function has been extensively studied in neurons, where it contributes to several fundamental processes such as neuronal differentiation *(50)*, excitability *(51)* and synaptic plasticity *(52)*, mostly by mediating BDNF stimuli through its interaction with Trk and p75 receptors *(36, 53, 54)*. In the absence of Kidins220, the expression of both TrkB isoforms is reduced in embryonic as well as in postnatal astrocyte cultures. In embryonic cultures, Kidins220^−/−^ astrocytes are characterized by reduced sustained activation of the MAPK pathway suggesting that similar to neurons *(36, 37)*, Kidins220 modulates MAPK activation downstream of full-length TrkB in astrocytes. In postnatal astrocytes, Kidins220 contributes to both full-length and truncated TrkB-induced [Ca^2+^]_i_ signaling, as both components are reduced in Kidins220-depleted cells. Of note, the interaction between Kidins220 and TrkB involves the transmembrane domain of the receptor *(36)* that is shared between the two receptor isoforms. Our gene expression analysis revealed reduced expression of Kir4.1 in Kidins220-depleted astrocytes. Interestingly, similar results have been recently described in cultured astrocytes lacking TrkB-T1 *(9–12)*, further supporting the notion that Kidins220 and TrkB receptors are part of the same signaling system. Together with the increased expression of the transient receptor potential cation channel subfamily V member 4 (TRPV4) that we previously described *(33)*, this observation suggests that the deficiency of Kidins220 subjects astrocytes to a global stress reaction and contributes to explain the developmental and activity impairments observed in wild type neurons co-cultured with Kidins220^−/−^ cells *(33)*.

Overall, our work contributes to define the complex and debated mechanisms of neurotrophin signaling in astrocytes, showing that - at least *in vitro* - both full-length and truncated TrkB receptors are competent to activate distinct intracellular signaling pathways at distinct phases of astrocyte development. Furthermore, we identified Kidins220 as one of the cellular components endowing full-length and truncated TrkB receptor their signaling specificity. Our future studies will address how these findings will translate *in vivo,* under physiological and pathological conditions. In recent years, mutations in the *KIDINS220* gene have been associated with severe neurodevelopmental pathologies, whose main symptoms are intellectual disability and spastic paraplegia *(55–61)*. A better understanding of the physiological functions of neurotrophins and Kidins220 in neurons and glial cells is instrumental to address the molecular pathways leading to this severe neurological disorder, which are currently completely unknown.

## MATERIALS AND METHODS

### Animals

All embryos used in this study were obtained from crosses of Kidins220^+/-^ mice *(37, 52)* in the C57BL/6 background. Mice were mated overnight and separated in the morning. The development of the embryos was timed from the detection of a vaginal plug, which was considered day 0.5. Postnatal cultures were prepared from P0-P1 pups obtained from crosses of Kidins220^+/lox^ mice on the C57BL/6 background. All experiments were carried out in accordance with the guidelines established by the European Community Council (Directive 2010/63/EU of 22 September 2010) and were approved by the Italian Ministry of Health.

### Antibodies

The following primary antibodies were used: rabbit polyclonal anti-Kidins220 (GSC16, #AB34790, Abcam), rabbit polyclonal anti-TrkB (#07-225, Millipore), rabbit monoclonal anti-phosphorylated TrkB (Tyr516, #4619, Cell Signaling) rabbit monoclonal anti-GAPDH (14C10, #2118, Cell signaling), rabbit monoclonal anti-phosphorylated MAPK1/2 (Thr202/Tyr204, #4377, Cell signaling), rabbit polyclonal anti-MAPK1/2 (#06-182, Millipore), rabbit monoclonal anti-phosphorylated Akt (Ser473, #4058, Cell signaling), rabbit polyclonal anti-Akt (#9272, Cell signaling), rabbit polyclonal anti-phosphorylated PLCγ (Tyr783, #2821, Cell signaling), rabbit polyclonal anti-PLCγ (#2822, Cell signaling), mouse monoclonal anti-glial fibrillary acidic protein (GFAP, #G3893, Sigma-Aldrich).

Secondary antibodies for western blot analysis were ECL Plex goat anti-rabbit IgG-Cy5 (PA45012, GE Healthcare), ECL Plex goat anti-mouse IgG-Cy3 (PA43009, GE Healthcare), HRP-conjugated goat anti-rabbit antibodies (#31460, ThermoFisher scientific) and HRP-conjugated goat anti-mouse antibodies (#31430, ThermoFisher scientific). Fluorescently conjugated secondary antibodies for immunocytochemistry were from Molecular Probes (Thermo-Fisher Scientific; Alexa Fluor 488, #A11029; Alexa Fluor 647, #A21450). Hoechst (#B2261, Sigma-Aldrich) was used to stain nuclei.

### Primary astrocyte culture

E18.5 or P0-P1 cortices were dissected in ice-cold PBS, incubated with trypsin 0.25% and 1 mg/ml DNase I for 30 min at 37 °C, and mechanically dissociated. Cells were then re-suspended and plated on poly-D-lysine-coated flasks or glass coverslips, in MEM medium containing 10% FBS, 2 mM glutamine, 33 mM glucose and antibiotics (astrocyte culture medium). After 24 h, the medium was removed and replaced with fresh culture medium. After one week, half of the medium was replaced with fresh culture medium. Embryonic and postnatal cultures were composed of about 90% astrocytes and were totally devoid of neurons *(33)*.

### Lentivirus production and infection procedures

HEK293T cells were maintained in Iscove’s Modified Dulbecco’s Medium supplemented with 10% FBS, 2 mM Glutamine, 100 U/ml penicillin and 0.1 mg/ml streptomycin in a 5% CO_2_ humidified incubator at 37 °C. Cells were transfected with the Δ8.9 encapsidation plasmid, the VSVG envelope plasmid and the pLenti-PGK-Cre-EGFP or pLenti-PGK-ΔCre-EGFP plasmids *(40)* using the calcium phosphate method. The transfection medium was replaced by fresh medium after 16 h. Supernatants were collected 36 to 48 h after transfection, centrifuged to remove cell debris, passed through a 0.45 μm filter and ultracentrifuged 2 h at 20,000 x *g* at 4°C. Viral pellets were re-suspended in PBS, aliquoted and stored at −80 °C until use.

Confluent postnatal cultures were trypsinized and seeded on 6-well plates or glass coverslips coated with poly-D-lysine for subsequent experiments. Cells were infected 3 days later with lentiviruses encoding catalytically dead (ΔCre) or active Cre recombinase with the lowest infectious dose capable of transducing ≥95% of cells (dilution range 1:500 to 1:800) and used for experiments ≥7 days after transduction.

### Calcium imaging

Astrocytes at 3 DIV or at 7-8 DIV after lentivirus infection were loaded with 1 μg/ml Fura-2-AM (#F1221, ThermoFisher) in astrocyte culture medium for 30 min at 37 °C. Subsequently, cells were washed in recording buffer (10 mM HEPES pH 7.4, 150 mM NaCl, 3 mM KCl, 1 mM MgCl_2_, 10 mM Glucose and 2 mM CaCl_2_) 30 min at 37 °C to allow hydrolysis of the esterified groups. Coverslips with cells were mounted on the imaging chamber and loaded with 0.45 ml of recording buffer. Fura-2-loaded cultures were observed with an inverted Leica 6000 microscope using a HCX PL APO lambda blue 63.0×1.40 oil-immersion objective. For analyzing BDNF-evoked Ca^2+^ transients, 50 μl of BDNF solution (final concentration 20 ng/ml, as reported in *(7)*, #B3795, Sigma) were manually added to the culture medium 15 s after the beginning of the recordings. Where indicated, cells were preincubated with 150 nM K-252a (tyrosine kinase blocker, #K1639, Sigma), 1 μM ANA-12 (TrkB receptor antagonist, #SML0209, Sigma), 10 μM U73122 (PLCγ inhibitor, #U6756, Sigma) or 10 μM U73343 (inactive analog, #U6881, Sigma) for 10 min before the beginning of the recordings. Samples were excited at 340 and 380 nm and images of fluorescence emission at 510 nm were acquired using a Hamamatsu-C9100-02-LNK00 camera. Calcium levels were estimated from background subtracted ratio images (340/380nm) of Fura-2-loaded astrocytes at the cell body level according to the equation of Grynkiewicz *(62)*:

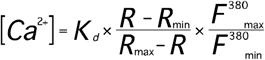

where R is the measured 340/380 nm ratio; R_mm_ and R_max_ are the ratios in the absence of Ca^2+^ or when Fura-2 is saturated by Ca^2+^, and F^380^_max_ and F^380^_min_ are the fluorescence intensity of 380 nm excitation at 0 Ca^2+^ and at Ca^2+^ saturation. To determine the Kd in our system, *in situ* calibration was performed by using 1 μM of the Ca^2+^-ionophore ionomycin in recording buffer containing an increasing concentration of Ca^2+^, ranging from 1 nM to 10 mM. [Ca^2+^]_i_ transients were defined for R/Ro ≥ 0.025.

### Biochemical techniques

Cells were washed once in ice-cold PBS and lysed in RIPA buffer (50 mM Tris-HCl pH 7.4, 150 mM NaCl, 2 mM EDTA, 1% NP40, 0.1% SDS) plus protease and phosphatase inhibitors (complete EDTA-free protease inhibitors, Roche Diagnostic; serine/threonine phosphatase inhibitor and tyrosine phosphatase inhibitor, Sigma). After centrifugation at 16,000 x *g* for 15 min at 4 °C, protein concentration was quantified by using the BCA Protein Assay kit (ThermoFisher Scientific). SDS-PAGE and western blotting were performed by using precast 4-12% NuPAGE Novex Bis-Tris Gels (Invitrogen). After incubation with primary antibodies, membranes were incubated with fluorescently conjugated secondary antibodies and revealed by a Typhoon Variable Mode Imager (GE Healthcare) or with HRP-conjugated secondary antibodies and ECL Prime Western Blotting System (#RPN2106, GE Healthcare) and imaged using a ChemiDoc imaging system (Biorad). Immunoreactive bands were quantified by using the ImageJ software. The fold change activation was calculated by first making the ratio of the intensity of phosphorylated bands to the total amount of corresponding protein for each sample, and then by dividing the phospho:total ratio of the treated samples by the phospho:total ratio of untreated control samples.

### Immunocytochemistry and image analysis

Cells were fixed in 4% PFA for 15 min, then washed in PBS. Cells were subsequently permeabilized with 0.2% Triton-X 100 in PBS for 10 min at room temperature (RT) then incubated with primary antibodies diluted in PBS 1% BSA overnight at 4°C or 2 h at RT. After washes in PBS, cells were incubated with fluorescent secondary antibodies diluted in PBS 1% BSA. After washes, coverslips were mounted with Mowiol.

Confocal stacks were acquired with a Leica SP8 using a 40X oil immersion objective (NA 1.40), with 1 μm between optical sections. Images were analyzed using the ImageJ software. The maximal fluorescence intensities of in-focus stacks were z-projected and the resulting images were automatically thresholded. Regions of interests (ROIs) were manually drawn around the borders of GFAP-positive cells in the red channel and fluorescence intensity was measured in the green channel (P-TrkB). Quantification was performed on 18-20 fields per condition from three independent cell preparations. Data are reported as the average P-TrkB fluorescence intensity of all GFAP+ cells in each field.

### Chronic BDNF treatment, RNA extraction and RT-qPCR

Three days after lentiviral infection, Kidins220^lox/lox^ astrocytes cultures were serum starved overnight then treated with 1 ng/ml BDNF on the following morning or left untreated. Treatment was repeated on the next two days and total RNA was extracted the following day with the QIAzol lysis reagent (Qiagen). The corresponding cDNAs were prepared by reverse transcription of 1 μg of RNA using the SuperScript III First-Strand Synthesis System (Invitrogen) with an oligo-dT primer according to the manufacturer’s instructions. The resulting cDNAs were used as a template for RT-qPCR using a CFX96 Real-Time PCR Detection System (Biorad) with a SYBR Green master mix (Qiagen). Thermal cycling parameters were 5 min at 95 °C, followed by 40 cycles of 95 °C for 15 s, and 60 °C for 45 s. The relative quantification in gene expression was determined using the ΔΔCt method. Data were normalized to Transferrin receptor protein 1 (TRFR), TATA-box-binding protein (TBP), and Tubulin beta-2A (TUBB2) by the multiple internal control gene method with GeNorm algorithm *(63)*. Sequences of the primers used are listed in **Supplementary Table 1**.

### Statistical analysis

Data are presented as means ± S.E.M. throughout the text. The distribution of the data was assessed using the D’Agostino-Pearson omnibus normality test. When comparing two groups unpaired twosided Student’s *t*-test and one-sample *t*-test were used, and equality of variances tested through the F test. When more than two groups were compared, two-way ANOVA followed by the Tukey’s post hoc multiple comparison test was performed to assess significance as indicated in figure legends, and equality of variances tested through the Brown-Forsythe’s and Bartlett’s test. Alpha levels for all tests were 0.05% (95% confidence intervals). No statistical methods were used to predetermine sample sizes, however sample sizes in this work (indicated in figure legends) are similar to those previously reported in the literature for similar experiments. The ROUT method with Q = 1% was used to identify outliers for exclusion from analysis. All statistical procedures were performed using GraphPad Prism 7 software (GraphPad Software, Inc).

## List of Supplementary Materials

**Supplementary Figure 1.** Activation of signaling pathways upon administration of 1 ng/ml BDNF in embryonic astrocytes.

**Supplementary Figure 2.** Activation of signaling pathways upon administration of 1 ng/ml BDNF in wild type postnatal astrocytes.

**Supplementary Figure 3.** Activation of signaling pathways upon administration of 1 ng/ml BDNF in Kidins220-deficient postnatal astrocytes.

**Supplementary Table 1.** List of primers used for the RT-qPCR analysis.

## Acknowledgements

We kindly acknowledge: drs. A. Mehilli, D. Moruzzo, and R. Navone for help in the maintenance, breeding and genotyping of the Kidins220 mouse strains; drs. R. Ciancio and I. Dallorto for the administrative support.

## Author contributions

F.J. performed the biochemical analysis and Ca^2+^ imaging experiments in embryonic astrocytes, part of the biochemical analysis and of the Ca^2+^ imaging experiments in postnatal astrocytes; M.A. performed immunocytochemistry in embryonic astrocytes, part of the biochemical analysis and of the Ca^2+^ imaging experiments in postnatal astrocytes, gene expression analysis; S.F. contributed to the designing of experiments and to discuss the results; F.B. contributed to the designing of experiments, to discuss the results and supported the project; F.C. designed the project, supervised all the experiments and discussed the results. All author contributed to writing the paper.

## Data and materials availability

The datasets used and/or analyzed during the current study are available from the corresponding author on reasonable request.

