## Supplementary material for "A developmental stage- and Kidins220/ARMS-dependent switch in astrocyte responsiveness to brain-derived neurotrophic factor"

**SUPPLEMENTARY FIGURE LEGENDS**

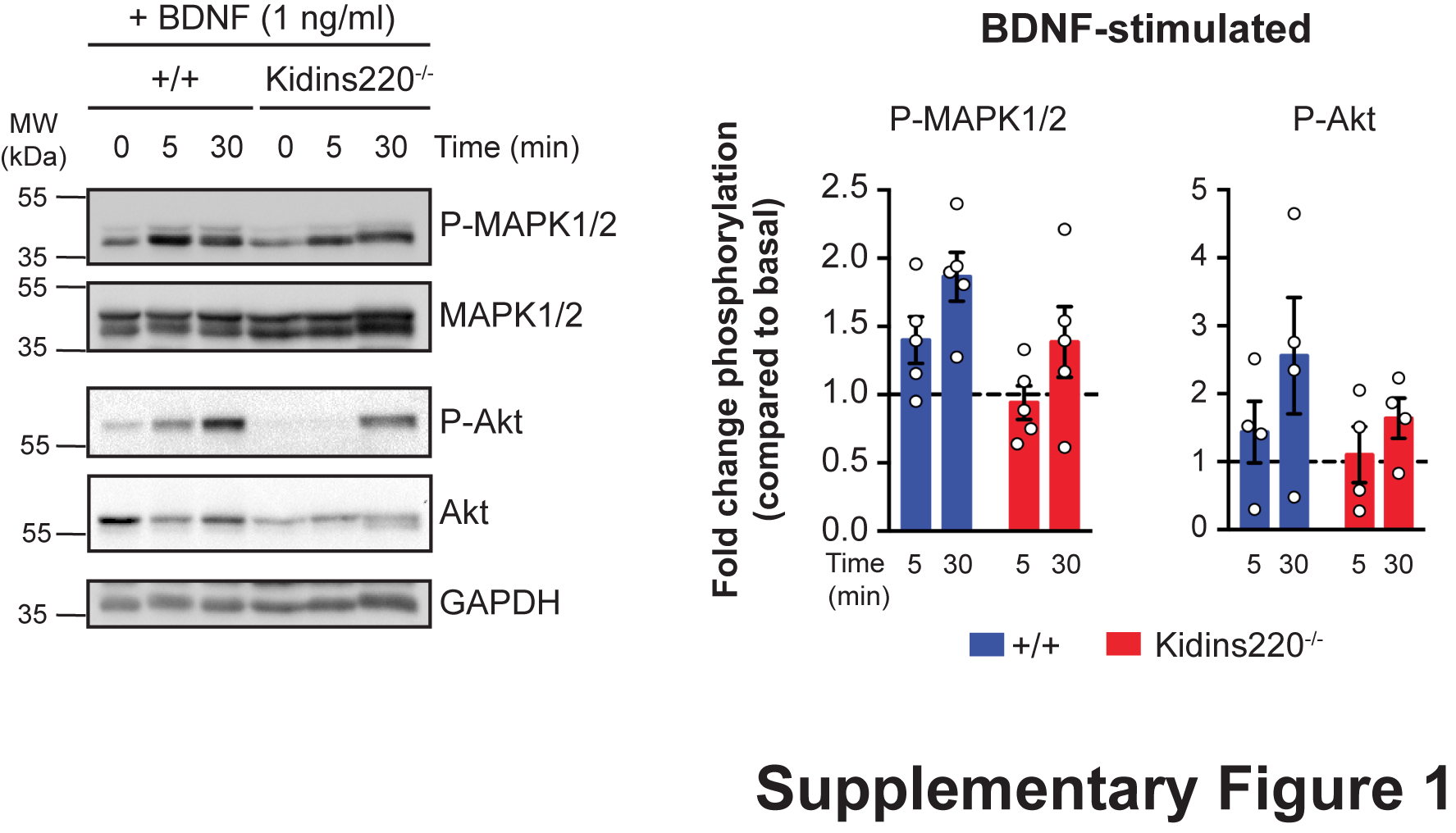

**Supplementary Figure 1. Activation of signaling pathways upon administration of 1 ng/ml BDNF in embryonic astrocytes.**

Wild type and Kidins220^-/-^ embryonic astrocyte cultures were treated with 1 ng/ml BDNF for 5 and 30 min or left untreated (time 0). Lysates were analyzed for phosphorylated MAPK1/2 (Thr202/Tyr204) and Akt (Ser473). PLCγ did not show any reliable activation at this BDNF concentration (not shown). Membranes were subsequently stripped and re-probed for the total amount of the same protein. *Left*: Representative immunoblots. *Right*: Time dependence of MAPK1/2 and Akt phosphorylation upon BDNF stimulation in wild type and Kidins220^-/-^ astrocytes. The graphs express the fold change activation of MAPK1/2 and Akt compared to the untreated phosphorylation levels for each genotype, set to 1 (dashed line in all graphs). For further details, see legend to Fig. 1 and Methods. For MAPK, we report the sum of MAPK1 and MAPK2 immunoreactivity. p>0.05, unpaired Student’s *t*-test, n=4-5 for both wild type and Kidins220^-/-^ cultures. Values are expressed as means ± S.E.M.

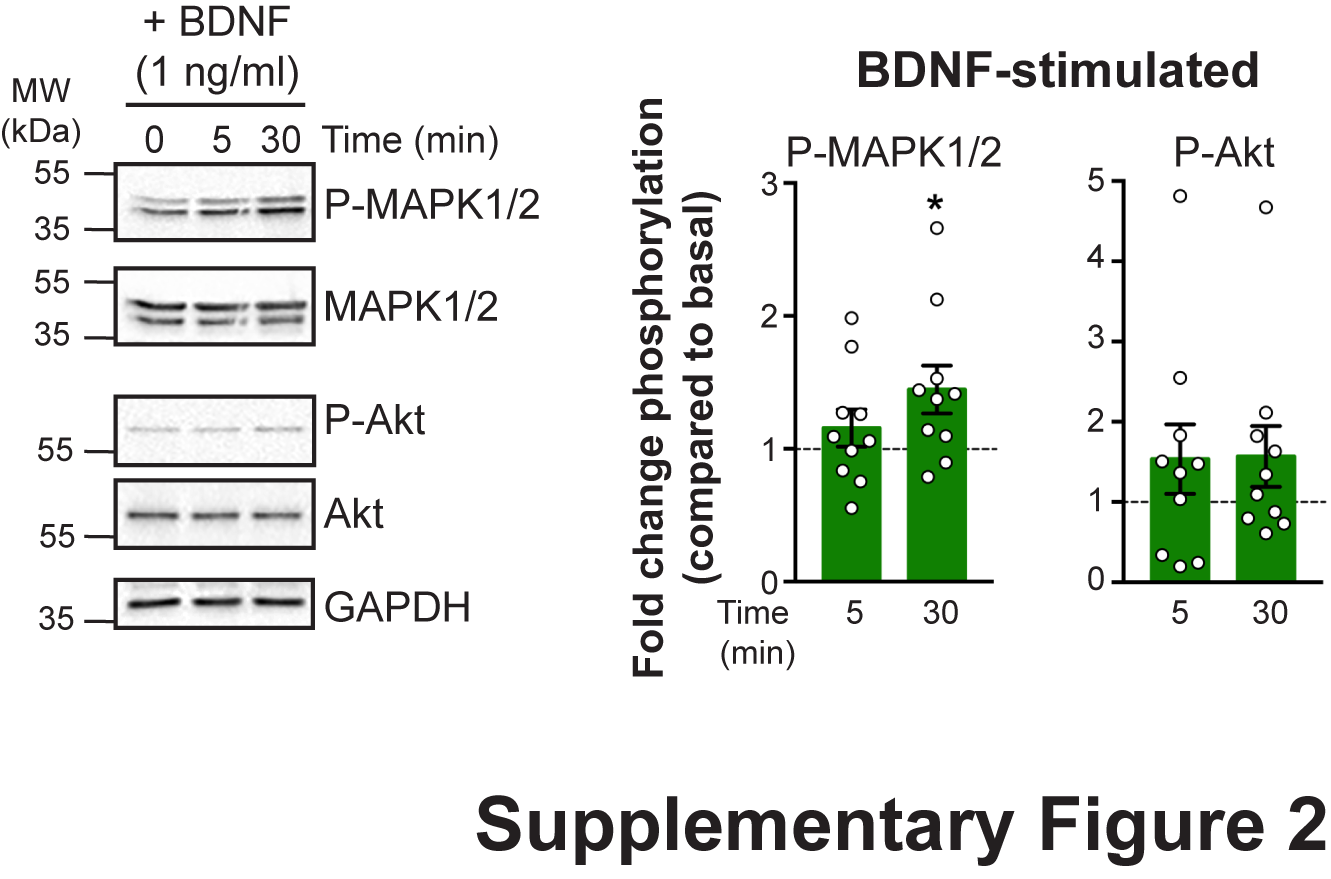

**Supplementary Figure 2. Activation of signaling pathways upon administration of 1 ng/ml BDNF in wild type postnatal astrocytes.**

Experiments as in Suppl. Figure 1, but for wild type postnatal astrocytes cultures. *p<0.05, one sample Student’s *t*-test compared to baseline, n=10 independent cultures. Values are expressed as means ± S.E.M.

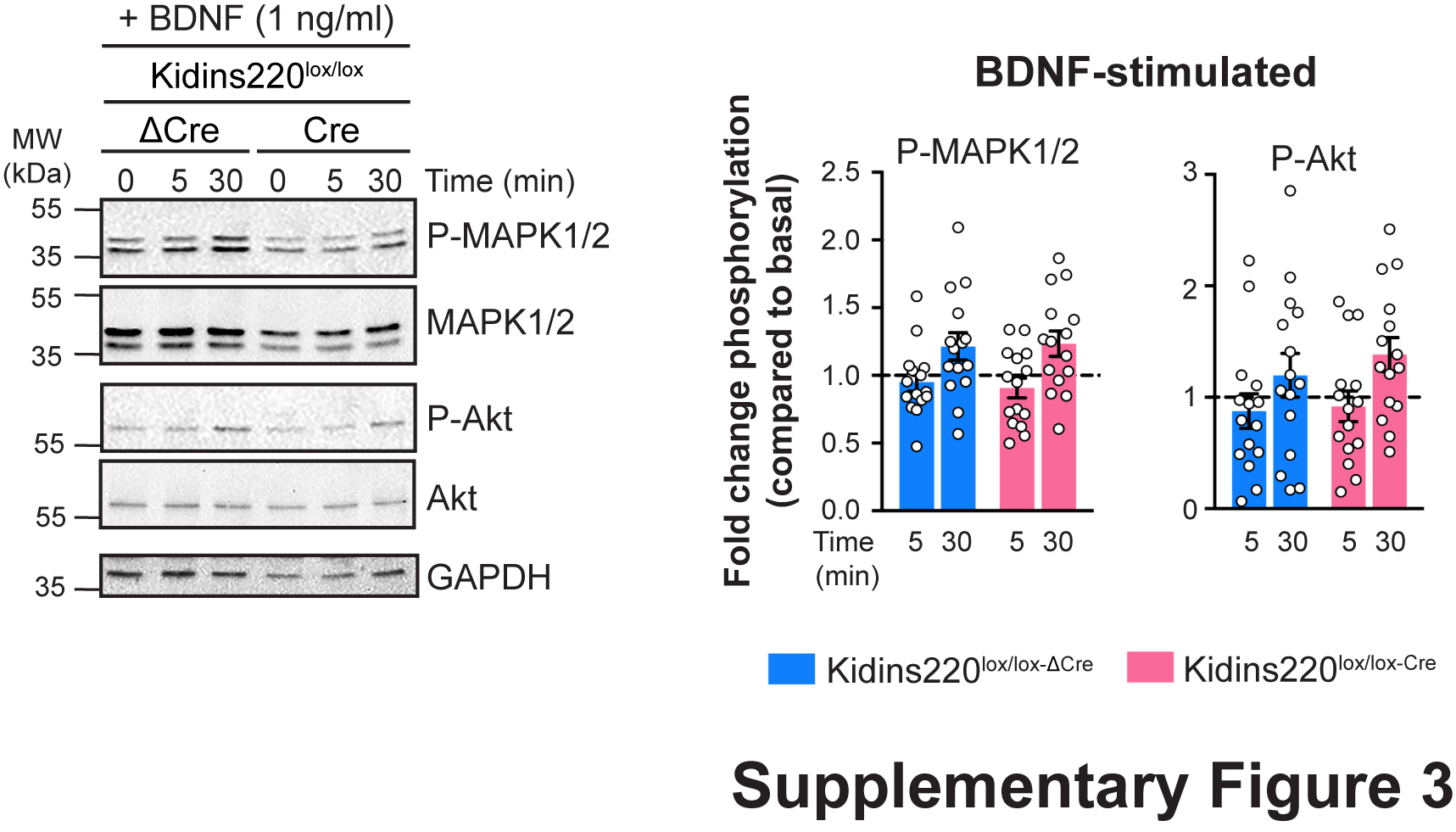

**Supplementary Figure 3. Activation of signaling pathways upon administration of 1 ng/ml BDNF in Kidins220-deficient postnatal astrocytes.**

Experiments as in Suppl. Figure 1, but for Kidins220^lox/lox-Cre^ and Kidins220^lox/lox-ΔCre^ astrocytes cultures. p>0.05, unpaired Student’s *t*-test, n=15 independent cultures. In all experiments, GAPDH was used as a loading control. Values are expressed as means ± S.E.M.

**SUPPLEMENTARY TABLE**

| **Gene** | **GenBank Accession** | **Forward primer (5’ 🡪 3’)** | **Reverse primer (5’ 🡪 3’)** |
| --- | --- | --- | --- |
| Slc1a2 (GLT-1)  Kcnj10 (Kir4.1)  Aqp4  Gja1 (Cnx43)  **Housekeeping genes**  TBP  TRFR  TUBB2 | [NM_001077514.4](https://www.ncbi.nlm.nih.gov/nucleotide/NM_001077514.4?report=genbank&log$=nucltop&blast_rank=5&RID=K2NURD91014)  NM_001039484.1  NM_009700.3  NM_010288.3  NM_013684.3  NM_011638.4  NM_009450.2 | ACTGGCTGCTGGATAGAATGA  GCCCCGCGATTTATCAGAG  CTGTGGCAGCGAGATAATGG  CTTTGACTTCAGCCTCCAAGG  ACTTCGTGCAAGAAATGCTGAAT  AGACCTTGCACTGTTTGGACATG  CAAGGCTTTCCTGCACTGGT | AATGGTGTCCAGCTCAGACT  TCCATTCTCACATTGCTCCG  GCCTTTCTGGGAACTCACAC  GGGCACCTCTCTTTCACTTAAT  CAGTTGTCCGTGGCTCTCTTATT  GGTGTGTATGGATCACCAGTTCCTA  AACTCCATCTCGTCCATGCC |

**Supplementary Table 1. List of primers used for the RT-qPCR analysis.**
